# *Lotus japonicus* CLV1-Like Receptor HAR1 Promotes Nitrogen Utilization and Growth Under Non-Symbiotic Conditions

**DOI:** 10.64898/2026.07.27.740878

**Authors:** Nao Okuma, Daisuke Sugiura, Ichiro Terashima, Masayoshi Kawaguchi

## Abstract

Legumes establish mutualistic symbiosis with nitrogen (N)-fixing bacteria, which allows them to utilize atmospheric N_2_. Because the maintenance of symbiosis requires abundant carbon (C) sources, legumes regulate the balance between carbon consumption and nitrogen acquisition by systemically controlling the nodule number through CLAVATA1 (CLV1)-like receptors. In *Lotus japonicus*, the CLV1-like receptor HYPERNODULATION ABERRANT ROOT FORMATION1 (HAR1) acts in shoots to regulate root nodulation and contributes to symbiotic C/N coordination. This raises the possibility that HAR1 may also influence plant growth and nitrogen utilization beyond symbiotic nodulation. In this study, we showed that HAR1 plays a critical role in regulating nitrogen use to enhance growth under conditions of high nitrate availability, even in non-symbiotic environments. Unlike the wild-type, the *har1* mutant failed to increase its growth in response to higher nitrate availability. This lack of growth response was associated with a lower rate of net biomass production per unit leaf area and a reduced capacity for biomass production per unit plant nitrogen. We further found that nitrate-responsive TCA cycle-related organic acids were higher in *har1* leaves than in wild-type leaves even under low nitrate conditions. Because the HAR1 mutation did not affect photosynthetic traits, we propose that HAR1 promotes growth under non-symbiotic conditions by coordinating nitrogen utilization with primary metabolism.

**One-sentence summary:** *Lotus japonicus* CLV1-like receptor HAR1 improves nitrogen utilization and growth at high nitrate conditions by regulating organic acid metabolism, independent of root nodule symbiosis.

## Introduction

Leguminous plants can utilize atmospheric nitrogen (N_2_) because they have evolved the ability of symbiotic interactions with soil nitrogen (N)-fixing bacteria, collectively referred to as rhizobia (Suzaki et al., 2015; Oldroyd and Leyser, 2020). In this symbiosis, rhizobia fix atmospheric N_2_ via bacterial nitrogenase activity and supply the host plant with ammonium. In return, plants supply rhizobia with photosynthates. Because photosynthate consumption for rhizobial symbiosis is very costly, legumes restrict nodule formation using a long-distance signaling mechanism referred to as autoregulation of nodulation (AON). The leucine-rich repeat receptor-like kinases (LRR-RLKs) HYPERNODULATION ABERRANT ROOT1 (HAR1), SUPER NUMERIC NODULES (SUNN), and NODULE AUTOREGULATION RECEPTOR KINASE (NARK), play central roles in AON in *Lotus japonicus*, *Medicago truncatula*, and soybean (*Glycine max*), respectively (Wopereis et al., 2000; Krusell et al., 2002; Nishimura et al., 2002; Searle et al., 2003; Schnabel et al., 2005). These LRR-RLKs perceive root-derived CLV3/ESR-related (CLE) signal peptides, which are synthesized in response to rhizobial infection or high soil nitrate levels, with the latter condition allowing plants to obtain sufficient nitrogen without relying on nitrogen-fixing root nodule symbiosis (Okamoto et al., 2009; Mortier et al., 2010; Reid et al., 2011; Okamoto et al., 2013; Reid et al., 2013; Nishida et al., 2016). After CLEs have been perceived by LRR-RLKs, a shoot-derived signal microRNA (miR2111) and cytokinin are translocated to roots and control nodulation in a manner dependent on the root-acting inhibitor, TOO MUCH LOVE (TML) F-box protein (Magori et al., 2009; Takahara et al., 2013; Sasaki et al., 2014; Tsikou et al., 2018; Gautrat et al., 2019; Gautrat et al., 2020; Okuma et al., 2020; Zhang et al., 2021). LRR-RLK mutants show impaired growth due to excessive nodule formation. Therefore, LRR-RLKs are critical for maintaining the proper balance between nitrogen acquisition and carbon supply in root nodule symbiosis through long-distance signaling.

*HAR1*, *SUNN*, and *NARK* are orthologues of Arabidopsis (*Arabidopsis thaliana*) *CLAVATA1* (*CLV1*), rice (*Oryza sativa*) *FLORAL ORGAN NUMBER1* (*FON1*), and tomato (*Solanum lycopersicum*) *FASCIATED AND BRANCHED* (*FAB*) that control shoot apical meristem (SAM) size (Nagasawa et al., 1996; Clark et al., 1997; Suzaki et al., 2004; Xu et al., 2015). Arabidopsis *clv1*, rice *fon1*, and tomato *fab* mutants show enlarged SAM size, whereas mutants of *L. japonicus har1*, *M. truncatula sunn*, and soybean *nark* exhibit normal SAM development (Wopereis et al., 2000; Nishimura et al., 2002; Searle et al., 2003; Schnabel et al., 2005). Therefore, the functions of CLV1-like LRR-RLKs are diversified between leguminous and non-leguminous plants. Under symbiotic conditions, HAR1, SUNN, and NARK regulate the symbiotic C/N balance by systemically controlling the number of root nodules through long-distance signaling that involves root–shoot–root communication.

However, it is unclear whether HAR1, SUNN, and NARK also contribute to the maintenance of C/N balance under non-symbiotic conditions. *L. japonicus har1*, *M. truncatula sunn*, and soybean *nark* exhibit pleiotropic phenotypes that are not directly related to root nodule symbiosis. For example, mutants of *L. japonicus har1* and soybean *nark* exhibit smaller leaf sizes and leaf cell numbers compared with wild-type under both symbiotic and non-symbiotic conditions (Ito et al., 2008; Tanabata et al., 2013). Roots of *L. japonicus har1*, *M. truncatula sunn*, and soybean *nark* mutants are shorter than wild-type roots, regardless of nodulation (Wopereis et al., 2000; Searle et al., 2003; Schnabel et al., 2005). Therefore, these LRR-RLKs may regulate growth and development in a root nodule symbiosis-independent manner.

In this study, we focused on growth suppression in the *L. japonicus har1* mutant. To address the unknown role of HAR1 in growth under non-symbiotic conditions, we investigated growth rates, photosynthetic traits, and amino acid and organic acid metabolism in the *har1* mutant under conditions with various nitrate levels. We found that HAR1 plays an important role in improving the efficiency of nitrogen utilization for growth. Growth inhibition in *har1* was particularly evident under high soil nitrate availability, and nitrogen productivity, a parameter representing the efficiency of nitrogen utilization for growth, was reduced in *har1*. This reduction was not accompanied by impaired photosynthetic nitrogen use.

Instead, *har1* reproducibly showed accumulation of TCA cycle-related organic acids compared with the wild-type. These observations suggest that HAR1 contributes to efficient nitrate-dependent growth under non-symbiotic conditions, potentially through the coordination of nitrogen utilization and primary metabolism.

## Results

### *har1* showed stunted growth under high nitrate conditions

*har1-7* is a null allele of *har1* with a non-sense mutation (W348stop) in the LRR domain. Here, we compared the growth of wild-type and *har1-7* mutant plants without rhizobial infection. One-week-old seedlings were transplanted into vermiculite-filled pots supplemented with Broughton and Dilworth solution (B&D) (Broughton and Dilworth 1971) containing either 1 mM or 5 mM nitrate. After 3 weeks, the shoot and root weights were measured. We found that *har1-7* exhibited a dwarf phenotype under 5 mM nitrate (Fig. 1a, b). The fresh weight (FW) of *har1-7* treated with 5 mM nitrate was significantly lower than that of the wild-type (Fig. 1c). The difference in dry weight (DW) under treatment with 5 mM nitrate showed a similar tendency to that for FW (*P* = 0.0502) (Fig. 1d), consistent with the finding for uninoculated soybean *nark* mutant (En6500) under higher nitrate concentrations (Takahashi et al., 1995). The shoot/root ratios (means ± SD) were 1.98 ± 0.53 in the wild-type and 2.33 ± 0.32 in *har1-7* at 1 mM nitrate conditions. At 5 mM nitrate conditions, the shoot/root ratios were 2.76 ± 0.42 in the wild-type and 2.73 ± 0.22 in *har1-7*. No significant differences were detected between genotypes at either nitrate condition (Welch’s *t*-test: 1 mM nitrate, *P* = 0.365; 5 mM nitrate, *P* = 0.904), indicating that plastic adjustment of the shoot/root ratio to nitrogen availability was normal in *har1-7*. These results suggest that HAR1 plays an important role in nitrate-dependent growth promotion.

**Figure 1:**
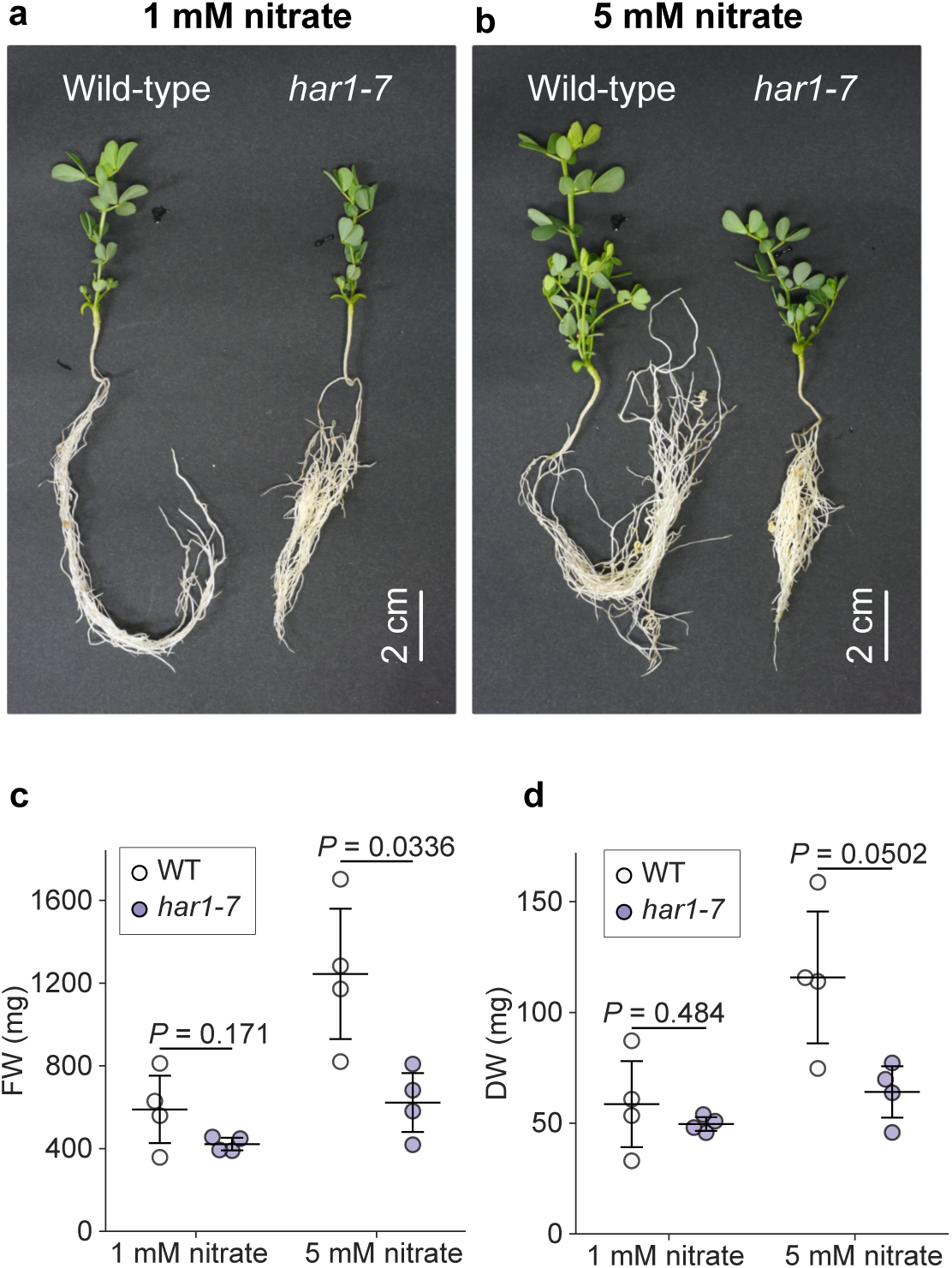
Phenotypic differences of wild-type and *har1-7* under different nitrate treatments. **a, b** Phenotypes of the wild-type and *har1-7* mutant grown on vermiculite supplemented with B&D containing 1 mM nitrate (a) or 5 mM nitrate (b) at 28 days after germination. **c, d** Fresh weights (c) and dry weights (d) of the wild-type and *har1-7* under each condition. Scatterplots show individual biological replicates as dots (*n* = 4). Error bars indicate mean ± standard deviation. Two-sided Welch’s *t*-test was used to determine statistical differences compared with wild-type.

### *har1* mutant showed lower nitrogen productivity compared with the wild-type under high nitrate conditions

As *har1-7* growth was specifically delayed at 5 mM nitrate, we analyzed the growth response of *har1-7* to different nitrate levels in the absence of rhizobia. One-week-old seedlings were grown in autoclaved vermiculite with B&D containing 1 mM nitrate for 2 weeks, and some were sampled (first sampling). The remaining seedlings were then transferred to B&D containing 1 mM, 2 mM, or 5 mM nitrate for the evaluation of growth responses to various nitrate levels. Plants were sampled 11 days after the transfer (second sampling) (Fig. 2a). We measured the DW and nitrogen contents of wild-type and *har1-7* plants. The DW of the wild-type increased in response to increasing nitrate, whereas the nitrate level had little effect on *har1*-*7* growth (Fig. 2b). The relative growth rate (RGR) of *har1-7* was similar regardless of nitrate concentration (Table 1). However, the values of total nitrogen content (per unit dry weight) were not significantly different between the wild-type and *har1-7* (Fig. 2c), suggesting that the efficiency of nitrogen utilization for growth was hindered in *har1-7*. To further examine the relationship between plant nitrogen status and growth, we compared nitrogen productivity (NP), which represents the rate of biomass gain per unit plant nitrogen. NP was calculated using measurements of dry weight and whole-plant nitrogen content obtained at the first and second harvests (see Materials and Methods for details). Indeed, NP was significantly decreased in *har1-7* under all conditions (Fig. 2d). Notably, the wild-type showed no significant difference in NP between the 1 and 2 mM nitrate conditions, whereas it was significantly lower under the 2 mM condition than under the 1 mM condition in *har1-7* (Fig. 2d). In the wild-type, a significant reduction in NP was observed only under the 5 mM nitrate condition, suggesting that the capacity for biomass production per unit plant nitrogen declined at a lower nitrate concentration in *har1-7*. The nitrogen content per leaf area was significantly higher in *har1-7* than in the wild-type at 2 mM nitrate (Fig. 2e).

**Figure 2:**
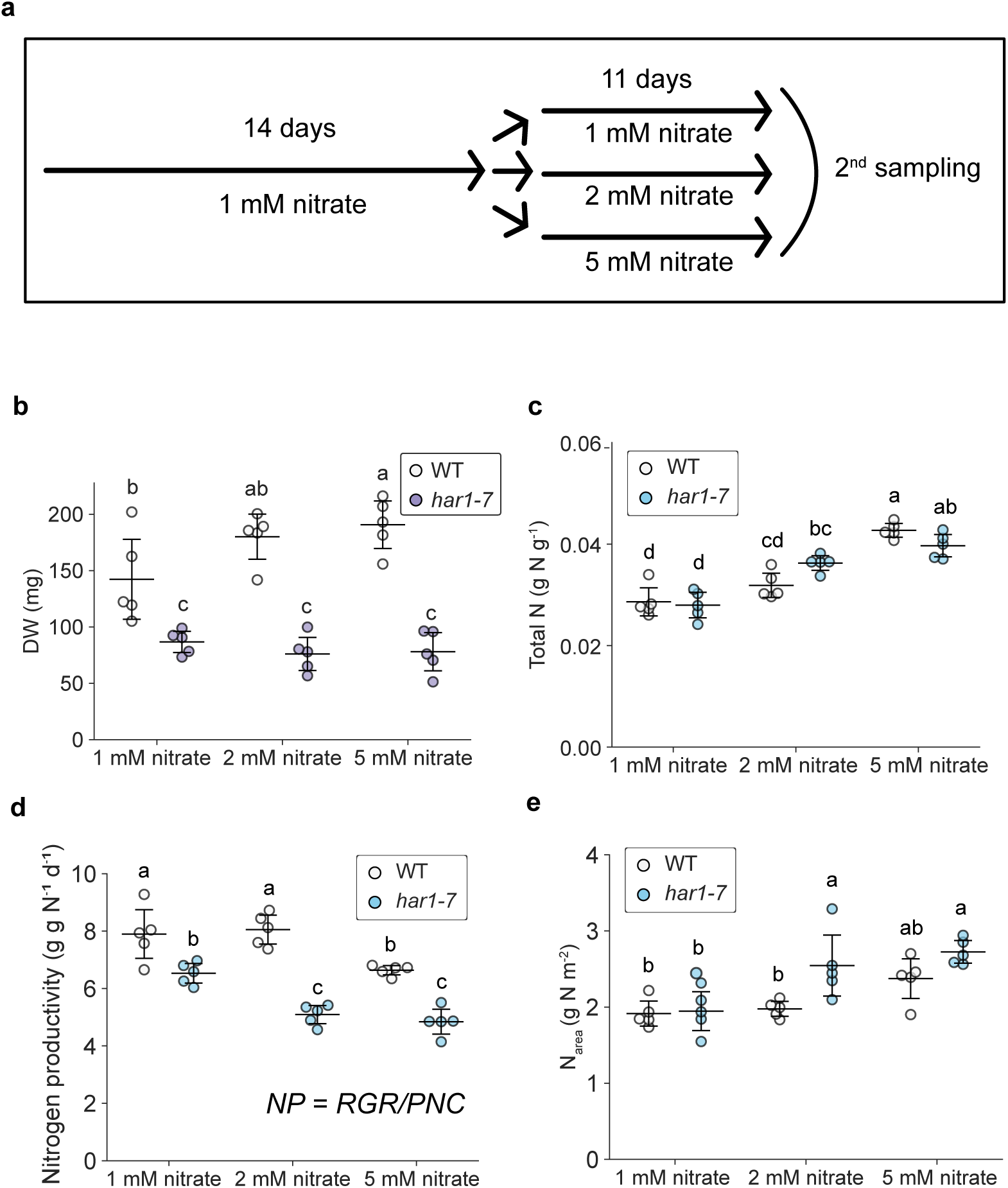
Comparison of the effects of nitrate concentration on growth rate and nitrogen contents of the wild-type and *har1-7*. **a** Schematic diagram to analyze growth rate in response to different levels of nitrate concentrations. **b** Dry weight of plants at second sampling. **c** Total nitrogen contents in plants at second sampling. **d** Nitrogen productivity (NP, g g N^−1^ d^−1^) of plants. NP is defined as relative growth rate (RGR) per plant nitrogen concentration (PNC, g N g^−1^), representing the nitrogen dependency of plant growth. **e** Nitrogen contents of wild-type and *har1-7* leaves (N_area_). Scatterplots show individual biological replicates as dots (*n* = 5). Error bars indicate mean ± standard deviation. Different letters indicate significant differences (*P* < 0.05) from Tukey’s honestly significant difference test.

**Table 1:**
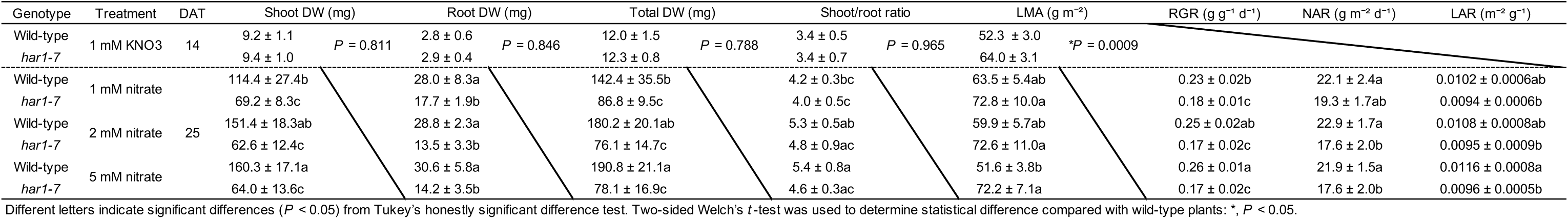
Dry weight (DW), shoot/root ratio, relative growth rate (RGR), net assimilation rate (NAR), and leaf area ratio (LAR). Values are shown as the mean ± standard deviation (*n* = 5). Fig. 2b-e were made from total DW, RGR, NAR, and LAR in this table, respectively. DAT, days after transplanting.

Furthermore, the net-assimilation rate (NAR, g m^−2^ day^−1^), representing the net-production efficiency of the leaves, was significantly lower in *har1-7* than in the wild-type specifically at 2 and 5 mM nitrate (Table 1). The leaf mass per area (LMA) of *har1-7* was also significantly higher at 5 mM nitrate (Table 1). These results suggested that although *har1-7* can accumulate nitrogen in the leaves, its net production efficiency is reduced. This lower efficiency is likely due to impairments in certain nitrogen utilization mechanisms within the leaves, particularly in the presence of 2 and 5 mM nitrate.

Additionally, the shoot nitrogen content per dry mass was lower, whereas the root nitrogen content per dry mass was higher in *har1-7* at 5 mM nitrate than in the wild-type (Supplementary Figure 1a,b). The nitrogen uptake rate per root dry mass (Osone et al., 2008) was significantly lower in *har1-7* than in the wild-type at 2 and 5 mM nitrate (Supplementary Figure 1c). These results suggest the involvement of HAR1 in both nitrogen allocation and uptake.

### The *har1* mutant can upregulate photosynthetic activity corresponding to the nitrogen level in leaves

Generally, leaf nitrogen contents are strongly correlated with the photosynthetic rate (Evans, 1989; Reich et al., 1995). We hypothesized that *har1-7* could not perform photosynthesis at the level implied by its leaf nitrogen content because it exhibited reduced RGR and NAR compared with the wild-type, particularly at higher nitrate levels, despite the relatively high level of nitrogen in its leaves (Fig. 2).

We evaluated the photosynthetic traits of *har1-7* after different nitrogen treatments. One-week-old seedlings were transplanted into vermiculite supplemented with B&D containing 2 mM nitrate and grown for 24 days. The nutrient medium was then changed to B&D containing either 2 mM nitrate (low-N) or 10 mM nitrate (high-N), and the physiological parameters were measured 3 days after the treatment (Experiment 1; Fig. 3a). We examined the photosynthetic rate. There were no significant differences in photosynthetic activity between high- and low-N conditions in either the wild-type or *har1-7* (Figure 3b, Table 2), suggesting that 2 mM nitrate was sufficient to maximize the photosynthetic rate under the present experimental conditions. Moreover, there were no differences in photosynthetic rate between wild-type and *har1-7* plants (Fig. 3b, Table 2). These results indicated that, in the presence of sufficient nitrate supply, the photosynthetic rate in *har1-7* was upregulated to the same extent as the rate in wild-type. The maximum rate of ribulose-1,5-bisphosphate (RuBP) carboxylation (*V*_cmax_) was proportional to leaf nitrogen content per area (*N*_area_) in all treatments (Supplementary Fig. 2a, Table 2).

**Figure 3:**
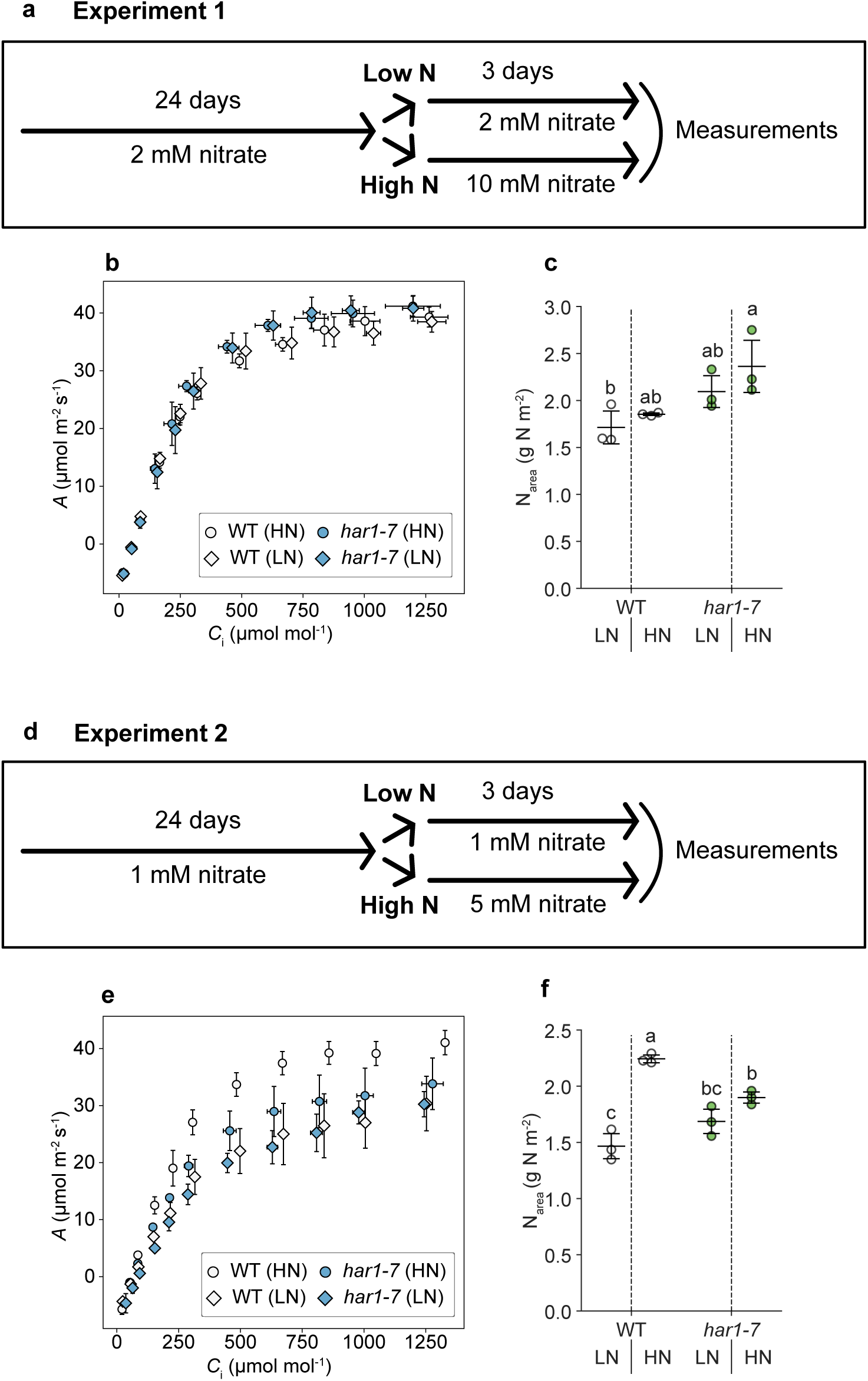
Photosynthetic trait of wild-type and *har1-7.* **a, d** Schematic diagram to analyze responses of CO_2_ assimilation to different nitrate concentrations. **b, e** Response of CO_2_ assimilation to intercellular CO_2_ concentration (*A*-*C*_i_ curve). Scatterplots show mean values. Values indicate mean ± standard deviation (*n =* 3). **c, f** Nitrogen contents of wild-type and *har1-7* leaves per area (*N*_area_) which was measured gas exchange. Scatterplots show individual biological replicates as dots. LN: low N; HN: high N. Error bars indicate mean ± standard deviation (*n =* 3). Different letters indicate significant differences (*P* < 0.05) from Tukey’s honestly significant difference test.

**Table 2:** Photosynthetic and physiological traits of wild-type and *har1-7*. The table of leaf nitrogen content per area (N_area_) and the maximum rate of ribulose-1,5-bisphosphate (RuBP) carboxylation rate (*V*_cmax_), the maximum electron transport rate *(J*_max_), leaf mass per area (LMA), and carbon content. Each value indicates the mean ± standard deviation (*n* = 3).

| Experiment | Genotype | Treatment | $N_{\text{area}}$ (g N m<sup>-2</sup>) | | $V_{\text{cmax}}$ (μmol m<sup>-2</sup> s<sup>-1</sup>) | | $J_{\text{max}}$ (μmol m<sup>-2</sup> s<sup>-1</sup>) | | LMA (g m<sup>-2</sup>) | | Carbon (%) | | C:N ratio | |
| --- | --- | --- | --- | --- | --- | --- | --- | --- | --- | --- | --- | --- | --- | --- |
| Experiment 1 | Wild-type | Low N (2 mM to 2 mM) | 1.72 ± 0.18b | $P = 0.318$ | 90.66 ± 9.36 | $P = 0.755$ | 179.81 ± 11.66 | $P = 0.799$ | 50.83 ± 4.80 | $P = 0.863$ | 42.44 ± 0.44 | $P = 0.351$ | 12.69 ± 1.57 | $P = 0.490$ |
| | | High N (2 mM to 10 mM) | 1.86 ± 0.02ab | | 88.40 ± 1.95 | | 182.57 ± 8.33 | | 51.62 ± 3.75 | | 42.07 ± 0.24 | | 11.72 ± 0.91 | |
| | *har1-7* | Low N (2 mM to 2 mM) | 2.10 ± 0.17ab | $P = 0.307$ | 91.81 ± 7.13 | $P = 0.112$ | 199.32 ± 11.01 | $P = 0.892$ | 57.26 ± 2.79 | $P = 0.366$ | 41.63 ± 0.69 | $P = 0.398$ | 11.51 ± 1.53 | $P = 0.595$ |
| | | High N (2 mM to 10 mM) | 2.37 ± 0.28a | | 104.54 ± 5.71 | | 200.58 ± 5.60 | | 61.67 ± 5.47 | | 40.82 ± 0.99 | | 10.74 ± 1.13 | |
| Experiment 2 | Wild-type | Low N (1 mM to 1 mM) | 1.47 ± 0.12c | $*P = 0.001$ | 63.17 ± 11.80 | $*P = 0.035$ | 139.71 ± 25.16 | $*P = 0.042$ | 103.33 ± 10.32 | $P = 0.111$ | 42.82 ± 0.66ab | $P = 0.080$ | 30.14 ± 1.40a | $*P = 0.003$ |
| | | High N (1 mM to 5 mM) | 2.25 ± 0.04a | | 94.15 ± 7.42 | | 196.41 ± 10.58 | | 75.90 ± 16.01 | | 44.31 ± 0.62a | | 15.00 ± 3.16b | |
| | *har1-7* | Low N (1 mM to 1 mM) | 1.69 ± 0.11bc | $P = 0.065$ | 63.14 ± 12.62 | $P = 0.350$ | 144.39 ± 9.22 | $P = 0.454$ | 104.22 ± 4.25 | $P = 0.710$ | 42.99 ± 0.43ab | $P = 0.339$ | 26.64 ± 1.51a | $P = 0.121$ |
| | | High N (1 mM to 5 mM) | 1.90 ± 0.05b | | 74.49 ± 8.43 | | 159.61 ± 24.27 | | 106.16 ± 5.40 | | 42.48 ± 0.52b | | 23.76 ± 1.43a | |

Different letters indicate significant differences ( $P < 0.05$ ) from Tukey's honestly significant difference test. Two-sided Welch's t-test was used to determine statistical difference between low N and high N conditions within each genotype: \*, $P < 0.05$ .

Under the present experimental conditions, the photosynthesis rate reached saturation at 2 mM nitrate in both the wild-type and *har1-7*; further increases in the nitrate level did not significantly alter photosynthetic activity.

Next, we evaluated the photosynthetic traits at lower nitrate levels. The nitrate level was set to 1 mM for the first 24 days of growth, after which the nutrient medium was changed to B&D containing either 1 mM nitrate (low-N) or 5 mM nitrate (high-N) (Experiment 2; Fig. 3d). In wild-type plants, both the photosynthetic rate and N_area_ were increased in response to high-N treatment, compared with low-N conditions (Figure 3e,f, Table 2). In contrast, no such increases in photosynthetic rate or N_area_ in response to high-N conditions were observed in *har1-7* (Fig. 3e,f, Table 2). The relationship between *V*_cmax_ and N_area_ was similar in wild-type and *har1-7* plants, as shown in Experiment 1 (Supplementary Figure 2b, Table 2). These results suggested that, despite the inability of *har1-7* to accumulate nitrogen in its leaves during short-term increases in nitrogen availability, *har1-7* remains capable of upregulating the photosynthetic rate consistent with its N_area_. Next, we performed two additional experiments with different nitrate conditions and plant ages (Experiments 3 and 4; see Materials and methods), and we analyzed relationships between N_area_ and the photosynthetic rate of *har1-7* using data from all four independent experiments (Supplementary Figure 2c, d). Analysis of covariance (ANCOVA) was used to assess the impact of N_area_ on the photosynthetic rate across wild-type and *har1-7*. The results confirmed significant positive correlations of N_area_ with *V*_cmax_ and the maximum electron transport rate (*J*_max_) (*V*_cmax_: *P =* 1.04e-14; *J*_max_: *P =* 2.97e-15); there was no significant effect of genotype (*V*_cmax_: *P =* 0.148; *J*_max_: *P =* 0.256) (Supplementary Figure 2c, d). Therefore, it is difficult to explain the stunted growth phenotype of *har1-7* under conditions of high nitrate availability based on the photosynthetic traits of *har1-7* in conditions of altered nitrate availability.

### *har1* mutant showed higher tricarboxylic acid (TCA) cycle-related organic acid levels in the leaves under high-nitrate conditions

*har1-7* was not able to utilize nitrogen for its growth well (Fig. 2), while nitrogen utilization in photosynthesis was not impaired in *har1-7* (Fig. 3, Supplementary Figure 2c,d). These results suggest that the growth phenotype of *har1-7* is not primarily caused by reduced photosynthetic nitrogen use. We therefore examined primary metabolism related to nitrogen and carbon utilization in *har1-7*, focusing on free amino acids and TCA cycle-related organic acids.

To examine possible functional sites of HAR1 involved in nitrogen metabolism, we first confirmed the spatial expression pattern of HAR1 under conditions of high nitrate availability. We generated *HAR1* promoter GUS transgenic lines (*proHAR1:GUS*). The *proHAR1:GUS* lines were stably transformed with the GUS reporter driven by a 2.5 kb DNA fragment upstream of the *HAR1* start codon. One-week-old seedlings of *proHAR1:GUS* lines were transplanted into vermiculite supplemented with B&D containing 5 mM nitrate and grown for 2 or 3 weeks.

Histochemical GUS staining assays were conducted using these transgenic plants. As previously reported, *HAR1* promoter activities were detectable in the vascular bundles of leaves and roots (Supplementary Figure 3a–d) (Nontachaiyapoom et al., 2007). In shoots, *HAR1* promoter activity was mainly detected in fully expanded leaves, but this activity was absent from younger leaves (Supplementary Figure 3e). This spatial expression pattern was similar to that of *NARK*, an orthologue of *HAR1* in soybean (Nontachaiyapoom et al., 2007). Therefore, we sampled the most recently fully expanded leaves of wild-type and *har1-7* treated with either 1 mM or 5 mM nitrate for further metabolic analyses.

We measured free amino acids and organic acids in leaves of the wild-type and *har1-7* in the absence of rhizobia using liquid chromatography–mass spectrometry (LC-MS). Youngest fully expanded leaves were collected from plants grown in four independent experiments under the same growth conditions. Amino acid profiles were first examined across four independent experimental batches (Fig. 4). Most amino acids did not show consistent genotype-dependent differences between the wild-type and *har1-7* under the same nitrate condition. Although several amino acids differed between 1 mM and 5 mM nitrate conditions, genotype-dependent changes in free amino acid levels were limited.

**Figure 4:**
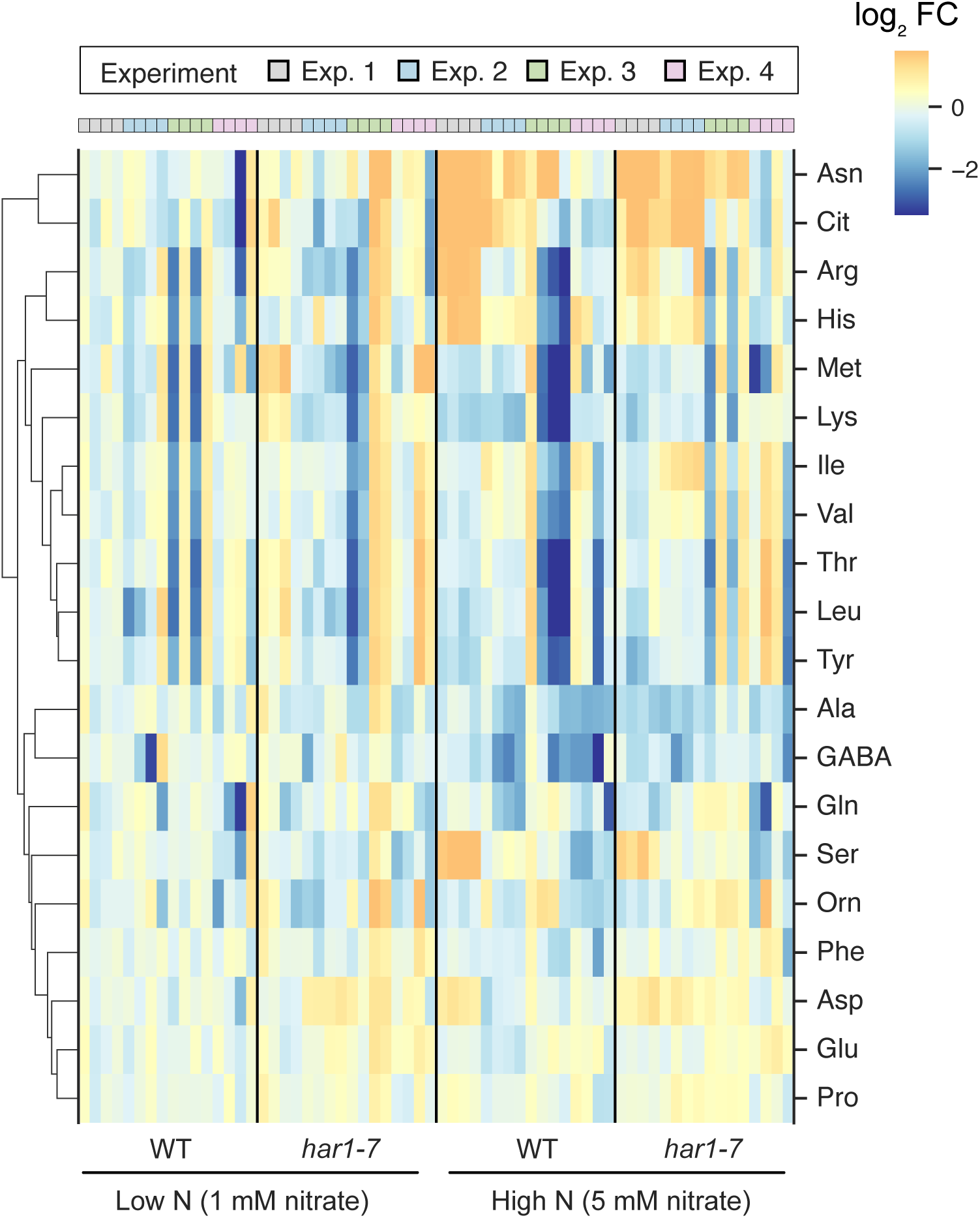
Amino acid profiles of wild-type and *har1-7* leaves. **a**, Heatmap showing amino acid levels in wild-type (WT) and *har1-7* leaves under 1 mM and 5 mM nitrate (Low-N and High-N) conditions across four independent experimental batches. Values are expressed as log fold changes (FC) relative to the mean value of WT under low-N conditions within each experimental batch. Each column represents an individual biological replicate. Columns are grouped by genotype–nitrate condition, and colored boxes above the heatmap indicate independent experimental batches. Hierarchical clustering of amino acids was performed based on Euclidean distance.

We next analyzed TCA cycle-related organic acids (Fig. 5, Supplementary Fig. 4). Among the organic acids examined, malic acid, citric acid, and succinic acid showed reproducible accumulation in *har1-7* across the four independent experiments (Fig. 5). These three organic acids were significantly higher in *har1-7* than in the wild-type under both 1 mM and 5 mM nitrate conditions. In addition, their levels tended to be higher under 5 mM nitrate conditions in both the wild-type and *har1-7,* suggesting that these organic acids respond to nitrate availability. These results indicate that *har1-7* shows altered accumulation of TCA cycle-related organic acids, whereas broad genotype-dependent changes in free amino acid contents were not observed.

**Figure 5:**
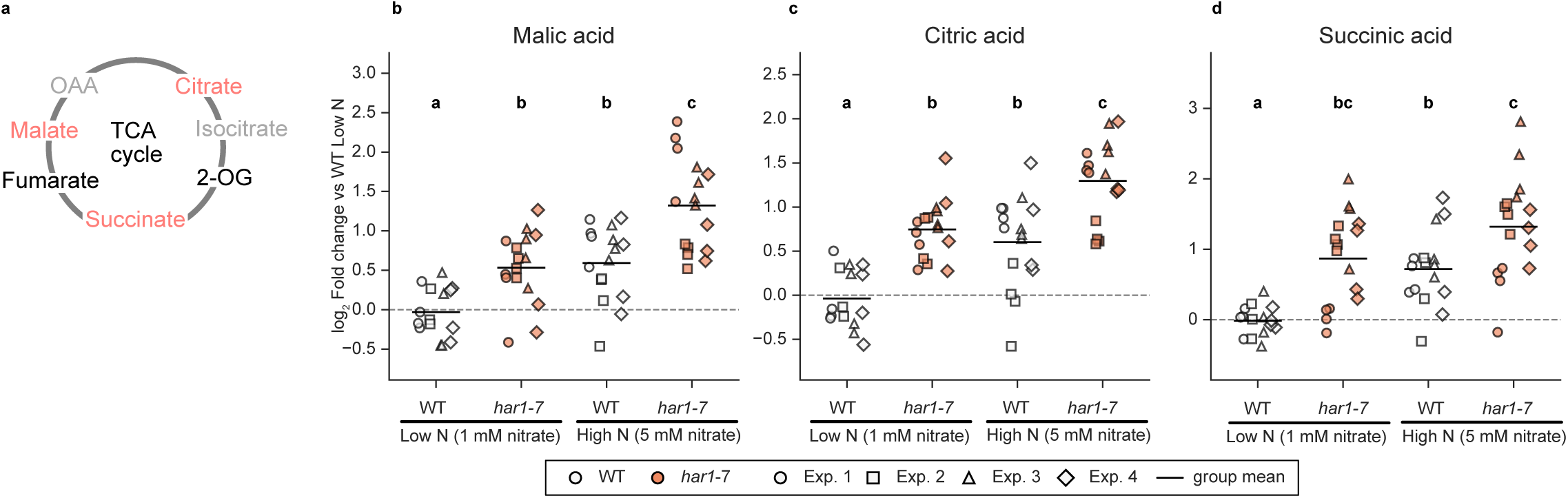
Accumulation of major organic acids in wild-type and har1-7 leaves. **a,** Schematic diagram of the tricarboxylic acid (TCA) cycle. Metabolite colors indicate the following: red, significantly increased in *har1-7*; black, no significant difference; grey, not detected. **b–d,** Relative levels of malic acid (b), citric acid (c), and succinic acid (d) in wild-type (WT) and *har1-7* leaves under 1 mM and 5 mM nitrate (Low-N and High-N) conditions. Values are expressed as log fold changes relative to the mean value of WT under low-N conditions within each experimental batch. Each dot represents an individual biological replicate; marker shapes indicate independent experimental batches. Black horizontal bars indicate group means. Different letters indicate significant differences among groups based on Tukey-adjusted pairwise comparisons of estimated marginal means from a linear model accounting for experimental batch effects (*P* < 0.05).

## Discussion

The balance of N and C contents is essential for precise plant growth and development (Nunes-Nesi et al., 2010). In leguminous plants, CLV1-like LRR-RLKs (e.g., HAR1, SUNN, and NARK) are presumed to play crucial roles in balancing symbiotic N acquisition and C consumption during root nodule symbiosis by restricting nodule numbers. Although *har1* mutants exhibit a pleiotropic phenotype even under non-symbiotic conditions, the functional significance of HAR1 beyond symbiosis has yet to be fully elucidated. In this study, we investigated the role of HAR1 in plant growth and nitrogen utilization under non-symbiotic conditions. We found that *har1* was unable to enhance growth in response to increasing nitrate availability (Figs. 1 and 2). This growth phenotype was accompanied by reduced nitrogen productivity, whereas photosynthetic nitrogen utilization was not impaired (Fig. 3; Supplementary Figure 2). These results suggest that the reduced growth of *har1-7* is unlikely to be primarily explained by impaired photosynthetic traits. Our metabolite analysis further showed that TCA cycle-related organic acids, particularly malic acid, citric acid, and succinic acid, reproducibly accumulated in *har1* leaves (Fig. 5, Supplementary Figure 4). In contrast, free amino acid contents in leaves did not show consistent genotype-dependent changes (Fig. 4). Because organic acids in the TCA cycle are closely linked to carbon metabolism and nitrogen assimilation, their accumulation may reflect an alteration in nitrate-associated primary metabolism in *har1*. Together, our results suggest that HAR1 contributes to efficient nitrate-dependent growth under non-symbiotic conditions, possibly by linking nitrogen utilization with primary carbon metabolism. The altered TCA cycle-related organic acid pools in *har1* provide a metabolic clue to this role, although further flux-based analyses will be required to clarify how HAR1 affects organic acid metabolism. Given that HAR1 orthologues are conserved in both nodulating and non-nodulating plants, this function may reflect a broader role of CLV1-like receptors beyond nodulation control.

We found that nitrogen productivity (NP) remained relatively stable in the wild-type between 1 and 2 mM nitrate conditions and declined only at 5 mM nitrate, whereas a reduction was already apparent in *har1-7* at 2 mM nitrate. This pattern suggests that the capacity for biomass production per unit plant nitrogen became limited at a lower nitrate concentration in *har1-7* (Fig. 2). The decline in NP observed in the wild-type at 5 mM nitrate further suggests that this concentration may exceed the range within which *L. japonicus* can efficiently use nitrogen for growth in our experimental conditions. We also showed that the HAR1 mutation did not significantly affect the relationship between photosynthetic rate and leaf N_area_ (Supplementary Fig. 2) or photosynthetic capacity under 2–10 mM nitrate conditions, in which nitrogen availability was sufficient to maintain photosynthesis (Fig. 3b).

These findings suggest that HAR1 may contribute to the utilization of nitrogen for biomass production rather than to the maintenance of photosynthetic capacity under nitrate-sufficient conditions. Instead, HAR1 may contribute to the effective allocation and utilization of nitrogen for plant growth under nitrate-sufficient conditions. We also found that *har1* could not adequately uptake nitrate according to root biomass at 2 and 5 mM nitrate compared with the wild-type (Supplementary Figure 1). Therefore, HAR1 may enhance nitrate uptake in response to nitrate in soil (Supplementary Figure 1). In contrast, the root nitrogen content per dry mass was significantly higher, whereas the shoot nitrogen content per dry mass was significantly lower, in *har1* than in the wild-type (Supplementary Figure 1). These differences were only observed with 5 mM nitrate treatment, which is probably an excess nitrate concentration for *L. japonicus* (see the 5 mM nitrate condition in Fig. 2d). Furthermore, unlike wild-type, *har1* did not exhibit a significant increase in leaf nitrogen content after 3 days of treatment with 5 mM nitrate (Fig. 3f). HAR1 likely facilitates immediate nitrogen transport from roots to the shoots in the presence of a high nitrate concentration.

Organic acids in the TCA cycle provide carbon skeletons for nitrogen assimilation and amino acid biosynthesis and therefore represent an important link between carbon metabolism, nitrogen metabolism, and growth (Scheible et al., 1997; Foyer et al., 2003; Sweetlove et al., 2010). Nitrate availability can induce coordinated changes in carbon and nitrogen metabolism, including the activation of organic acid metabolism and the accumulation of organic acids such as malate and citrate (Scheible et al., 1997). Consistent with this view, malic acid, citric acid, and succinic acid tended to accumulate to higher levels under 5 mM nitrate conditions in both the leaves of wild-type and *har1* in the present study. However, these organic acids were also reproducibly higher in leaves of *har1* than in the wild-type under both the low and high nitrate conditions. This pattern raises the possibility that HAR1 is involved in the regulation of TCA cycle-related organic acid pools in leaves during nitrate-dependent growth. In contrast to the reproducible changes in organic acids, free amino acid contents in leaves did not show broad or consistent genotype-dependent changes. Therefore, the reduced nitrogen productivity of *har1* may be more likely associated with altered organic acid accumulation than with broad changes in free amino acid pools. Because metabolite abundance represents pool size rather than metabolic flux (Heise et al., 2015), the increased levels of malic acid, citric acid, and succinic acid do not necessarily indicate enhanced TCA cycle activity. Nevertheless, together with the reduced nitrogen productivity of *har1*, this organic acid accumulation in *har1* leaves suggests a possible link between HAR1 function and nitrate-associated primary metabolism.

The LMA of *har1* was significantly higher than that of the wild-type under conditions of 2 and 5 mM nitrate (Table 1,2). Because the expansion of thinner leaves represents a major strategy for effective utilization of carbon and nitrogen under suitable light and soil nitrogen availability (Wright et al., 2004; Poorter et al., 2009), HAR1 may enhance effective leaf expansion via leaf morphological modifications when nitrogen is highly available. Moreover, the higher LMA of *har1* may have resulted from the accumulation of non-structural carbohydrates, such as soluble sugars and starch, although carbohydrate contents were not directly measured in the present study. The higher LMA, together with the lower leaf area ratio (LAR, m^-2^ g^-1^), indicates that *har1* had a smaller leaf area per unit plant biomass. This pattern was associated with lower NAR under 2 and 5 mM nitrate conditions (Table 1). Although carbohydrate accumulation has been associated with feedback regulation of photosynthesis in physiological and transcriptomic studies of legume species (Sugiura et al., 2019; Ozawa et al., 2023), our photosynthetic analyses did not indicate reduced photosynthetic capacity in *har1*. Instead, the accumulation of TCA cycle-related organic acids may represent another metabolic feature associated with inefficient conversion of available nitrogen and carbon resources into growth.

Together, these observations suggest that the reduced growth of *har1* may be associated with altered coordination among nitrogen utilization, primary metabolism, and leaf expansion. Further analyses of metabolic fluxes and carbohydrate accumulation will be required to clarify how HAR1 contributes to nitrate-dependent growth.

CLV1-like LRR-RLKs, such as HAR1, are well conserved in other plant species. Additionally, major components of AON (e.g., CLE peptide, miR2111, and TML) are conserved among various species. The functional significance of the CLE– CLV1-like LRR-RLK signaling module is well-characterized in Arabidopsis, rice, and tomato, particularly concerning the maintenance of shoot and floral apical meristems. However, functions involved in maintaining the meristematic region have not been observed in leguminous plants. Notably, the Arabidopsis CLE-CLV1 signaling module functions under nitrogen-deficient conditions; it negatively regulates expansion of the lateral root system in response to nitrogen deficiency (Araya et al., 2014). Moreover, miR2111, a downstream factor of leguminous CLV1-like LRR-RLKs, regulates lateral root emergence in *L. japonicus* and Arabidopsis in response to nitrate (Sexauer et al., 2023). Therefore, the function of the CLE-CLV1 signaling module conserved in legumes and Arabidopsis is closely related to the nitrogen response. Further investigations of the functional similarities and differences in AON-related genes within leguminous and non-leguminous plants are necessary to determine the evolutionary origins of systems involved in stabilizing root nodule symbiosis. Our findings provide insight into the possible importance of legume CLV1-like RLKs in C/N primary metabolism, particularly in terms of promoting growth under conditions of sufficient nitrate availability, regardless of root nodule symbiosis.

Additional studies are required to characterize how CLV1-like RLKs expressed in vascular bundles regulate the metabolism of key amino acids and organic acids in leaves.

## Materials and methods

### Plant materials

*Lotus japonicus* accession MG-20 was used as the wild-type in the present study (Kawaguchi, 2000). *har1-7*, a null allele of *har1* with a non-sense mutation (W348stop) in the LRR domain, was derived from MG-20 (Magori et al., 2009). proHAR1:GUS lines were generated on the MG-20 background.

### Plant growth conditions and sampling

Surface-sterilized seeds were germinated on 0.9% agar medium containing Broughton and Dilworth solution (B&D) without any nitrogen sources and grown for 1 week at 24°C (16 h light, 8 h dark, photosynthetic photon flux density: PPFD; about 55 μmol m^−2^ s^−1^) (Broughton and Dilworth, 1971). All seedlings used in this study were prepared under these conditions. KNO_3_ was used as the nitrate source in all experiments and was supplied at the indicated concentrations.

To determine the function of HAR1 under non-symbiotic conditions, we sterilized all instruments used in the experiments with 70% ethanol or by autoclaving to prevent rhizobial infection of *L. japonicus*. We confirmed the absence of nodules and nodule primordia on plant roots in all experiments. KNO was used as the nitrate source in all experiments and was supplied at the indicated concentrations.

For fresh weight and dry weight measurements (Fig.1 and Table 1) and the determination of metabolite amounts (Fig. 4, 5), 1-week-old seedlings of MG-20 and *har1-7* were transplanted into the pots (770 mL) filled with autoclaved vermiculite (PPFD; 250∼350 μmol m^−2^ s^−1^). The pots were placed on plastic trays (approximate height 6 cm and base 39 cm × 27 cm, 8 pots per tray), supplemented with approximately 700 mL of B&D containing 1 or 5 mM nitrate every 2 days. 21 days after transplanting, plants were measured for physiological and morphological parameters. For LC/MS analyses, the youngest fully expanded leaves were sampled and lyophilized using VD-800F (TAITEC). LC-MS analyses were performed using samples obtained from four independent experiments conducted under the same growth conditions.

For growth analyses, 1-week-old seedlings of MG-20 and *har1-7* were transplanted into pots (350 mL) filled with autoclaved vermiculite and grown in a growth chamber (24°C, 16 h light, 8 h dark, PPFD; about 500 μmol m^−2^ s^−1^). For each nitrate treatment, 12 plants were placed on the tray supplemented with approximately 1000 mL of B&D containing 1 mM nitrate every 2 days. The first sampling was conducted 14 days after transplanting. Four plants were sampled and divided into leaves, stems, and roots. After determination of the leaf area, the plants were oven-dried at 80°C and used for further analyses. The remaining plants were transferred to new plastic trays and supplemented with approximately 700 mL of B&D containing 1, 2, or 5 mM nitrate every 2 days. The second sampling was conducted 11 days after transfer in the same manner as the first sampling.

For photosynthesis measurements, 1-week-old seedlings of MG-20 and *har1-7* were transplanted into pots (770 mL) filled with autoclaved vermiculite. The pots were placed on a plastic tray (6 pots per tray) and grown in a growth chamber (24°C, 16 h light, 8 h dark, PPFD; about 500 μmol m^−2^ s^−1^). These trays were supplemented with approximately 500 mL of B&D containing the indicated concentration of nitrate every 2 days. In Experiments 1 and 2, the level of nitrate in B&D was modified to the concentration shown in the figures on day 24 (Fig. 3a, d). Photosynthetic traits were measured after 3 days (see below). In Experiment 3, 1-week-old seedlings were grown on vermiculite supplemented with B&D containing 1 mM nitrate for 40 days.

Then, the nutritional medium was changed to B&D containing 1 or 5 mM nitrate, and photosynthetic traits were measured 3 days later. In Experiment 4, 1-week-old seedlings were grown on vermiculite supplemented with B&D containing 5 mM nitrate for 35 days, and photosynthetic traits were measured. Leaves used for photosynthesis measurements were oven-dried at 80°C for further analyses.

Measurements and sampling were conducted in the same manner as Experiments 1 and 2. In Experiment 4, plants were supplied with B&D containing 1 mM nitrate and grown for 40 days. The plants with pots were placed on new trays and supplied with B&D containing either 1 or 5 mM nitrate. After 3 days, measurements and sampling were conducted in the same manner as the other experiments.

Two lines of *proHAR1:GUS* plants (Line #1 and Line #2) were used for promoter GUS assays under two conditions. In the first condition, 1-week-old seedlings of Line #1 and Line #2 were transplanted into autoclaved vermiculite-filled magenta boxes supplemented with B&D containing 5 mM nitrate (PPFD; about 500 μmol m^−2^ s^−1^). After 14 days, plants were sampled and used for GUS assays. In another experiment, 1-week-old seedlings of Line #1 were grown under the conditions described above for metabolite determination. These shoots were sampled and used for GUS assays.

### Plasmid construction and stable transformation

A 2.5 kb DNA fragment upstream of *HAR1* was amplified from *L. japonicus* ecotype MG-20 genomic DNA using Tks Gflex DNA Polymerase (Takara) with specific primers (5′-TCAGTCGACTGGATCAGTACGATTCATCGATGGTG-3′ and 5′- GTGCGGCCGCGAATTGCATTTGTTATGCTGAGTGT-3′). The amplified fragment was cloned into EcoRI- and BamHI-digested pENTR-1A (Invitrogen) using an In-Fusion HD Cloning Kit (Clontech). The insert was transferred into pMDC162 by LR clonase (Thermo Fisher Scientific) (Curtis and Grossniklaus, 2003). Stable transformation of *L. japonicus* MG-20 was performed using *Agrobacterium tumefaciens*-mediated method as previously described (Okuma et al., 2020).

### Growth analysis

The RGR (relative growth rate, g g^−1^ d^−1^) and NAR (net assimilation rate, g m^−2^ d^−1^) were determined from the first and the second sampling, assuming that changes in biomass and leaf area were exponential:

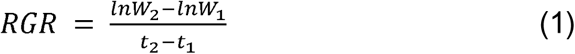

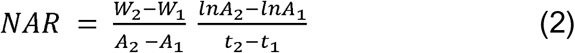

Where *W*_1,2_ and *A*_1,2_ are total biomass (g) and leaf area (m^2^) at the first sampling (*t*_1_, 14 days after transplanting) and the second sampling (*t*_2_, 25 days after transplanting), respectively. LAR (leaf area ratio, m^−2^ g^−1^) is further calculated from the following equation:

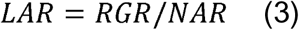

### Leaf gas exchange measurements

Gas exchange measurements were conducted using the youngest fully expanded leaves with a portable gas exchange system, LI-6800 (LI-COR Biosciences). The maximum carboxylation rate (*V*_cmax_, μmol m^□2^ s^□1^) and maximum electron transport rate (*J*_max_, μmol m^□2^ s^□1^) were determined from the CO_2_ response curves by fitting the following equations (Farquhar et al., 1980):

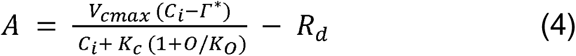

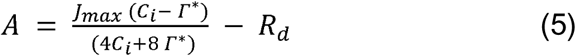

Where *A* is CO_2_ assimilation rate, *C_i_* is the intracellular CO_2_ concentration, *Γ*\* is the CO_2_ compensation point in the absence of photorespiration (36.9 μmol mol^□1^), *O* is the partial pressure of O_2_ (200 mmol mol^□1^), and *R_d_* is the day respiration. *K_c_* and *K_o_* are the Michaelis-Menten constants of Rubisco for CO_2_ (259 μmol mol^□1^) and O_2_ (179 mmol mol^□1^), respectively (von Caemmerer et al., 1994). The CO_2_ response curves were obtained by measuring *A* at ambient CO_2_ concentrations of 0, 50, 100, 200, 400, 600, 800, 1000, 1200, and 1500 μmol mol⁻¹. All measurements were conducted at PPFD of 1500 μmol m^-2^ s^-1^, leaf temperature of 25°C, and relative humidity of 50%. *V*_cmax_ and *J*_max_ were calculated using *C_i_*< 300 μmol mol^□1^ and > 300 μmol mol^□1^, respectively.

### Nitrogen analysis

Leaves used for the photosynthesis measurements were finely ground using a mill (TissueLyser II, Retsch, Haan, Germany), and leaf nitrogen content per area (N_area_, g N m^−2^) was determined with a CN analyzer (Vario EL III, Elementar, Germany).

Dried samples obtained in the growth analysis were also finely ground using the mill; the nitrogen concentration of each organ and the plant nitrogen concentration (PNC, g N g^−1^) were determined with the CN analyzer. Nitrogen productivity (NP, g g N^−1^ d^−1^), representing the nitrogen dependency of plant growth (Ågren, 1985), was defined as RGR per PNC, and was calculated assuming exponential growth of both dry weight and plant nitrogen content between the harvests as follows:

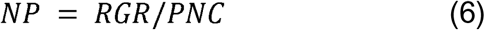

Nitrogen uptake rate per root dry mass (g N g^−1^ d^−1^) was determined from the first and the second sampling as follows:

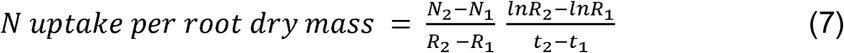

Where *N*_1,2_ and *N*_1,2_ are total plant nitrogen content (g N) and root mass (g) at the first sampling (*t*_1_, 14 days after transplanting) and the second sampling (*t*_2_, 25 days after transplanting), respectively.

### LC-MS

The lyophilized youngest fully expanded leaves of wild-type and *har1-7* plants, grown as described above, were used to determine amino acid and TCA cycle-related organic acid contents. Briefly, 400 μL of extraction solution (80% methanol, 0.1% formic acid, 10 pmol/μL^-1^ L-malic acid-^13^C_4_, and 1 pmol μL^-1^ ^13^C^15^N valine), which had been chilled to −20°C, was added to 2 mg of prepared leaves. The samples were then vortexed for 5 min and centrifuged at 13,000 rpm for 10 min at 4°C. Each supernatant (300 μL) was transferred to a new sample tube. The samples were then dried using a speed vacuum concentrator and reconstituted in 20 μL of 0.1% formic acid. Finally, 2 μL of each sample was analyzed by LC-MS (Triple TOF® 5600+ system SCIEX) combined with ACQUITY UPLC H-Class system (Waters).

Metabolites were separated using Discovery HS F5-3 column (2.1 mm ×150 mm, particle size 3 µm; Sigma) with a gradient elution of mobile phase A (0.1% formic acid in H_2_O) and mobile phase B (0.1% formic acid in acetonitrile) (0 min: 0% B; 2 min: 0% B; 5 min: 25% B; 11 min: 35% B; 15 min 95% B) at an eluent flow rate of 100 μL min^-1^ at room temperature. Amino acids were measured with positive-mode electrospray ionization with multiple-reaction monitoring (Phe: 166.1 to 120.1; Tyr: 182.1 to 136.1; Met: 150.1 to 56.06; Ile: 132.1 to 86.11; Leu: 132.1 to 86.11; Hyp: 132.1 to 86.08; Val: 118.1 to 55.06; ^13^C^15^N Val: 124.1 to 59.07; Glu: 148.1 to 84.06; Pro: 116.1 to 70.083; Asp: 134.2 to 74.042; Thr: 120.1 to 74.078; Ala: 90.1 to 44.065; Ser: 106.1 to 60.061; Lys: 147.1 to 84.101; Orn: 133.1 to 70.083; Asn: 133.1 to 74.042; His: 156.1 to 110.09; Arg: 175.1 to 70.083; GABA: 104.1 to 87.064; Cit: 176.1 to 70.083.). Organic acids were measured with negative-mode electrospray ionization with multiple-reaction monitoring (citric acid: 190.9 to 111.01; pyruvic acid: 86.9 to 87.01; succinate acid: 117 to 73.03; lactic acid: 88.9 to 88.9; malic acid: 133 to 133.02; glyoxylic acid: propionic acid: 73 to 45.006; □-ketoglutaric acid: 145 to 101.03; fumaric acid: 115 to 71.01; 4-methyl-2-ozovaleric: 129.1 to 69.04; ^13^C malic acid: 137 to 137.02).

### GUS staining assay

Before staining, shoots of *proHAR1:GUS* lines were incubated with ice-cold 90% acetone on ice for 10 min. Roots and acetone-treated shoots were incubated with GUS staining buffer (0.4 mg mL^-1^ X-Gluc, 50 mM phosphate buffer pH 7.0, 1 mM K_4_[Fe(CN)_6_], 1 mM K_3_[Fe(CN)_6_], and 0.1% Triton X-100) at 37°C for 18 h. Stained shoots were treated with acetic acid and ethanol (ratio 6:1) at 37°C for 1 h to remove the chlorophyll background. Stained shoots and roots were observed using an SZX16 stereomicroscope or a BX50 microscope (Olympus).

### Statistical analysis

Tukey’s honestly significant difference test (Tukey-HSD) was performed in R software (ver. 4.4.1). Two-sided Welch’s *t*-test and Pearson’s correlation coefficient analyses were performed in Python (ver. 3.12.13) with the SciPy library (ver. 1.11.3). Analysis of covariance (ANCOVA) was performed in Python using the Statsmodels library (ver. 0.14.6). For ANCOVA, photosynthetic parameters (*V*_cmax_ and *J*_max_) were analyzed using linear models of the form ‘*photosynthetic parameter*’ *∼ N_area_ + genotype,* where *N_area_* was included as a covariate and genotype was included as a categorical factor. For LC-MS metabolite data, relative peak areas from independent experimental batches were combined after sample labels were standardized. Four genotype–nitrate combinations were analyzed: WT under 1 mM nitrate; *har1-7* under 1 mM nitrate; WT under 5 mM nitrate; and *har1-7* under 5 mM nitrate. For each metabolite, natural-log-transformed relative peak areas were analyzed using a linear model of the form *log(relative area) ∼ ‘genotype–nitrate combination’ + experiment*. The experiment term was included to account for differences among independent LC-MS batches. Estimated marginal means were calculated from the fitted model, and pairwise comparisons among the four genotype–nitrate combinations were performed with Tukey adjustment using the emmeans package (ver. 2.0.2) in R. For visualization, metabolite levels were expressed as log□ fold changes relative to the mean value of wild-type under 1 mM nitrate conditions within each experimental batch.

## Acknowledgments

We thank Tomoko Mori for support for the LC-MS analyses; Takashi Soyano, Kensuke Kawade, and Taro Maeda for expert advice regarding experiments; Akiko Oda and Sachiko Tanaka for assistance with experiments; and the Trans-Omics Facility and Model Organisms Facility Center (Trans-Scale Biology Center, National Institute for Basic Biology) for technical support.

## Author contributions

N.O. and M. K. conceived the project and designed the experiments. D.S. and N.O. performed the photosynthetic and growth analyses. N.O. performed all other experiments. N.O. performed statistical analyses. N.O mainly wrote the paper, and all other authors contributed revisions. All authors approved the final version of this manuscript.

## Competing interests

The authors declare no competing interests.

Supplementary materials

## Supplementary materials

**Supplementary Figure 1: Nitrogen contents and uptake rates of wild-type and *har1-7* plants. a,b** Nitrogen contents of wild-type and *har1-7* shoots (shoot N) (a), and roots (root N) (b).

**c** Nitrogen uptake rate per root mass (g N g^−1^ d^−1^) of the wild-type and *har1-7*. Scatterplots show individual biological replicates as dots (*n* = 5). Error bars indicate mean ± standard deviation. Different letters indicate significant differences (*P* < 0.05) from Tukey’s honestly significant difference test.

**Supplementary Figure 2: Relationship between leaf nitrogen content and photosynthetic rate. a, b** Relationships between leaf nitrogen content per area (N_area_) and maximum rate of ribulose-1,5-bisphosphate (RuBP) carboxylation rate (*V*_cmax_) in Experiment 1 (a) and Experiment 2 (b). LN: low N; HN: high N. a; *R*^2^ = 0.318, b; *R*^2^ = 0.482.

**c, d** Relationship between the N_area_ and *V*_cmax_ (c) or the maximum electron transport rate (*J*_max_) (d) in all experiments (Experiments 1-4; see Materials and methods for details). c; *R*^2^ = 0.783, d; *R*^2^ = 0.803.

**Supplementary Figure 3: Detection of HAR1 promoter activities using promoter GUS assays. a–e** GUS expression controlled by a 2.5 kb DNA fragment upstream of the HAR1 coding region in shoots and roots of plants grown in closed magenta boxes (21 days after germination) (a–d) and shoots of plants grown in open pots (31 days after germination) (e). Two lines of transgenic plants, Line #1 (a–b, e) and Line #2 (c–d), showed similar GUS expression patterns. Black and white arrows indicate immature leaves where no GUS expression was observed.

**Supplementary Figure 4: Accumulation of organic acids in wild-type and *har1-7* leaves.** Relative levels of tricarboxylic acid (TCA) cycle-related organic acids in wild-type (WT) and *har1-7* leaves under low- andhigh-nitrate conditions. Values are expressed as log fold changes relative to the mean value of WT under low-N conditions within each experimental batch. Each dot represents an individual biological replicate; marker shapes indicate independent experimental batches. Black horizontal bars indicate group means. Different letters indicate significant differences among groups based on Tukey-adjusted pairwise comparisons of estimated marginal means from a linear model accounting for experimental batch effects (*P* < 0.05). Lactic acid and pyruvic acid were excluded from experiment 1 (Exp. 1), and fumaric acid was excluded from experiments 2 and 3 (Exp. 2 and 3), because distinct single peaks could not be reliably detected. The data for malic acid, citric acid, and succinic acid are identical to those presented in Fig. 5, but are shown here together with the other organic acids.

**Supplementary Table 1: Dry weight (DW) and fresh weight (FW) of plants under high- and low-nitrate treatments at 3 weeks after transplanting.** Values are shown as mean ± standard deviation (*n* = 4). Figure 1c and Figure 1d were made from total FW and total DW in this table, respectively.

