## Supplemental figures and tables for "*Lotus japonicus* CLV1-Like Receptor HAR1 Promotes Nitrogen Utilization and Growth Under Non-Symbiotic Conditions"

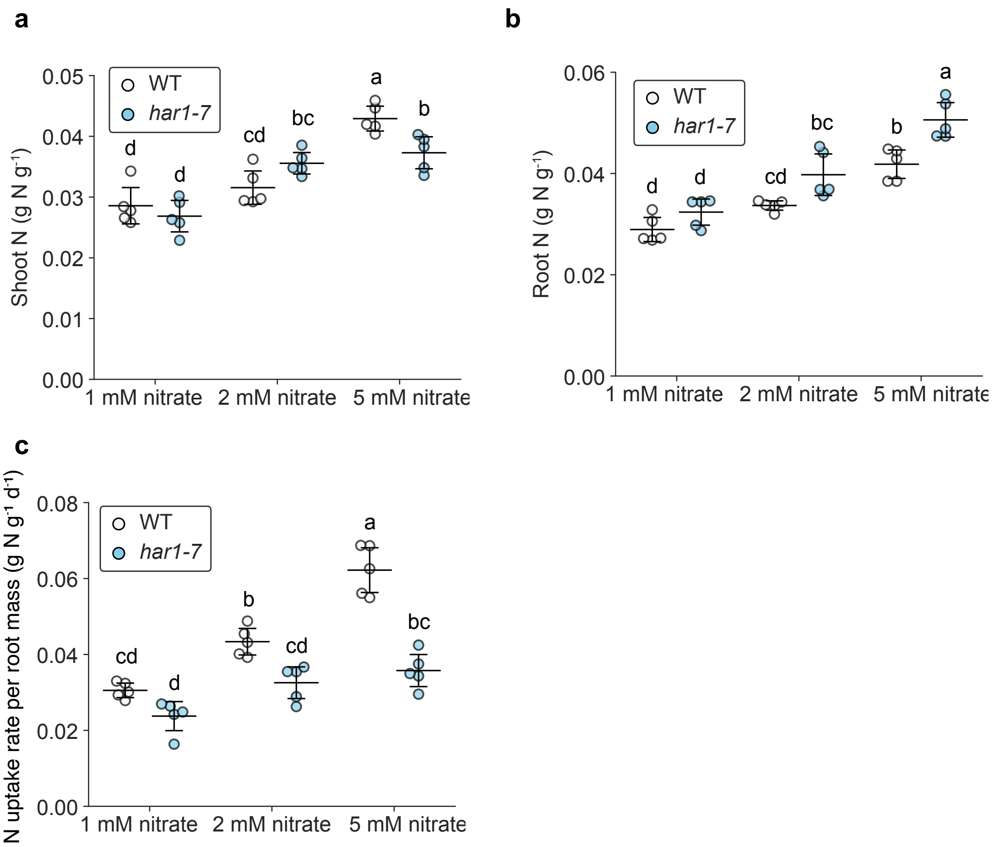


**Supplementary Figure 1: Nitrogen contents and uptake rates ofwild-type and *har1-7* plants.**

**a,b** Nitrogen contents of wild-type and *har1-7* shoots (shoot N) (a), and roots (root N) (b).

**c** Nitrogen uptake rate per root mass (g N $g^{-1}$ $d^{-1}$) of the wild-type and *har1-7*.

Scatterplots show individual biological replicates as dots ($n$ = 5). Error bars indicate mean ± standard deviation. Different letters indicate significant differences ($P < 0.05$) from Tukey's honestly significant difference test.

**Supplementary Table 1: Dry weight (DW) and the fresh weight (FW) of plants under high and low nitrate treatments at 3 weeks after transplanting.**
Values are shown as mean ± standard deviation (*n* = 4). Figure 1c and 1d were made from total FW and total DW in this table, respectively.

| Genotype | Treatment | Shoot DW (mg) | | Root DW (mg) | | Total DW (mg) | | Shoot/root ratio (DW) | | Shoot FW (mg) | | Root FW (mg) | | Total FW (mg) | | Shoot/root ratio (FW) | |
|---|---|---|---|---|---|---|---|---|---|---|---|---|---|---|---|---|---|
| Wild-type | 1 mM nitrate | 39.5 ± 16.3 | *P* = 0.647 | 19.2 ± 3.2 | *P* = 0.103 | 58.6 ± 19.4 | *P* = 0.484 | 2.0 ± 0.6 | *P* = 0.371 | 230.4 ± 89.3 | *P* = 0.433 | 359.1 ± 75.0 | *P* = 0.0633 | 589.5 ± 162.7 | *P* = 0.171 | 0.7 ± 0.2 | *P* = 0.140 |
| *har1-7* | | 34.7 ± 3.3 | | 15.0 ± 1.2 | | 49.7 ± 3.1 | | 2.4 ± 0.4 | | 183.5 ± 18.0 | | 238.2 ± 22.4 | | 421.7 ± 30.4 | | 0.8 ± 0.1 | |
| Wild-type | 5 mM nitrate | 84.6 ± 20.8 | **P* = 0.0439 | 31.3 ± 9.9 | *P* = 0.0867 | 115.8 ± 29.8 | *P* = 0.0502 | 2.8 ± 0.5 | *P* = 0.903 | 619.8 ± 142.6 | **P* = 0.0252 | 624.9 ± 182.2 | *P* = 0.0508 | 1244.7 ± 315.0 | **P* = 0.0336 | 1.1 ± 0.2 | *P* = 0.959 |
| *har1-7* | | 46.9 ± 8.7 | | 17.3 ± 3.3 | | 64.2 ± 11.7 | | 2.8 ± 0.3 | | 314.9 ± 82.4 | | 307.8 ± 62.7 | | 622.7 ± 142.8 | | 1.1 ± 0.1 | |

Two-sided Welch's *t*-test was used to determine statistical difference compared with wild-type plants: *, $P < 0.05$.

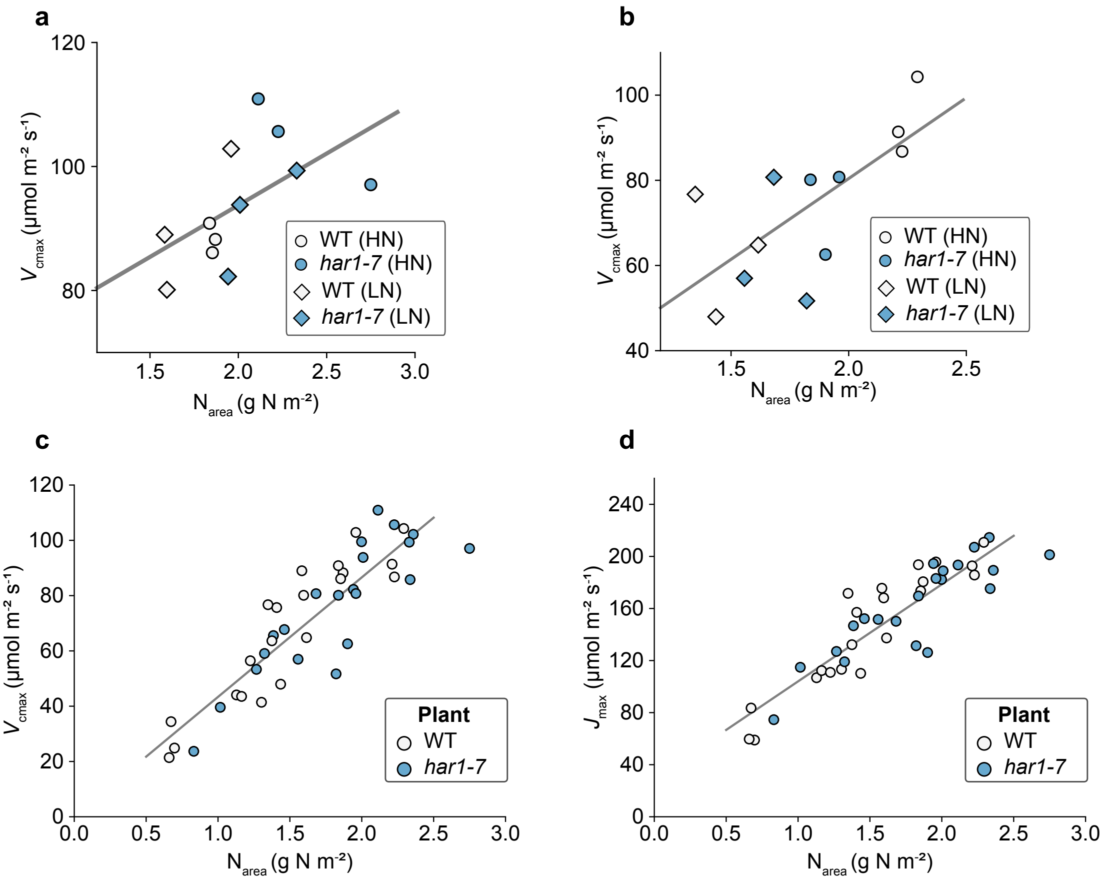


**Supplementary Figure 2: Relationship between leaf nitrogen content and photosynthetic rate.**

**a, b** Relationships between leaf nitrogen content per area ($N_{area}$) and maximum rate of ribulose-1,5-bisphosphate (RuBP) carboxylation rate ($V_{cmax}$) in Experiment 1 (a) and Experiment 2 (b). LN: low N; HN: high N.

a; $R^2$ = 0.318, b; $R^2$ = 0.482.

**c, d** Relationship between the $N_{area}$ and $V_{cmax}$ (c) or the maximum electron transport rate ($J_{max}$) (d) in all experiments (Experiments 1–4; see Materials and methods for details).

c; $R^2$ = 0.783, d; $R^2$ = 0.803.

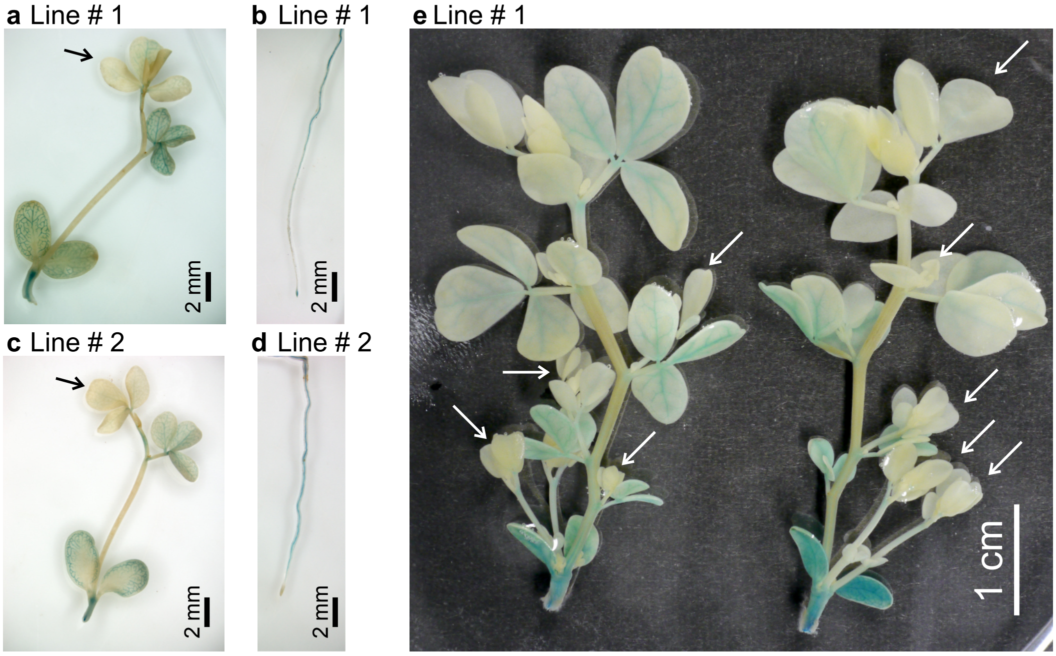


**Supplementary Figure 3: Detection of HAR1 promoter activities using promoter GUS assays.**
**a-e** GUS expression controlled by a 2.5 kb DNA fragment upstream of the HAR1 coding region in shoots and roots of plants grown in closed magenta boxes (21 days after germination) (a–d) and shoots of plants grown in opened pots (31 days after germination) (e).Two Lines of transgenic plants, line #1 (a-b, e) and Line #2 (c–d), showed similar GUS expression patterns. Black and white arrows indicate immature leaves where no GUS expression was observed.

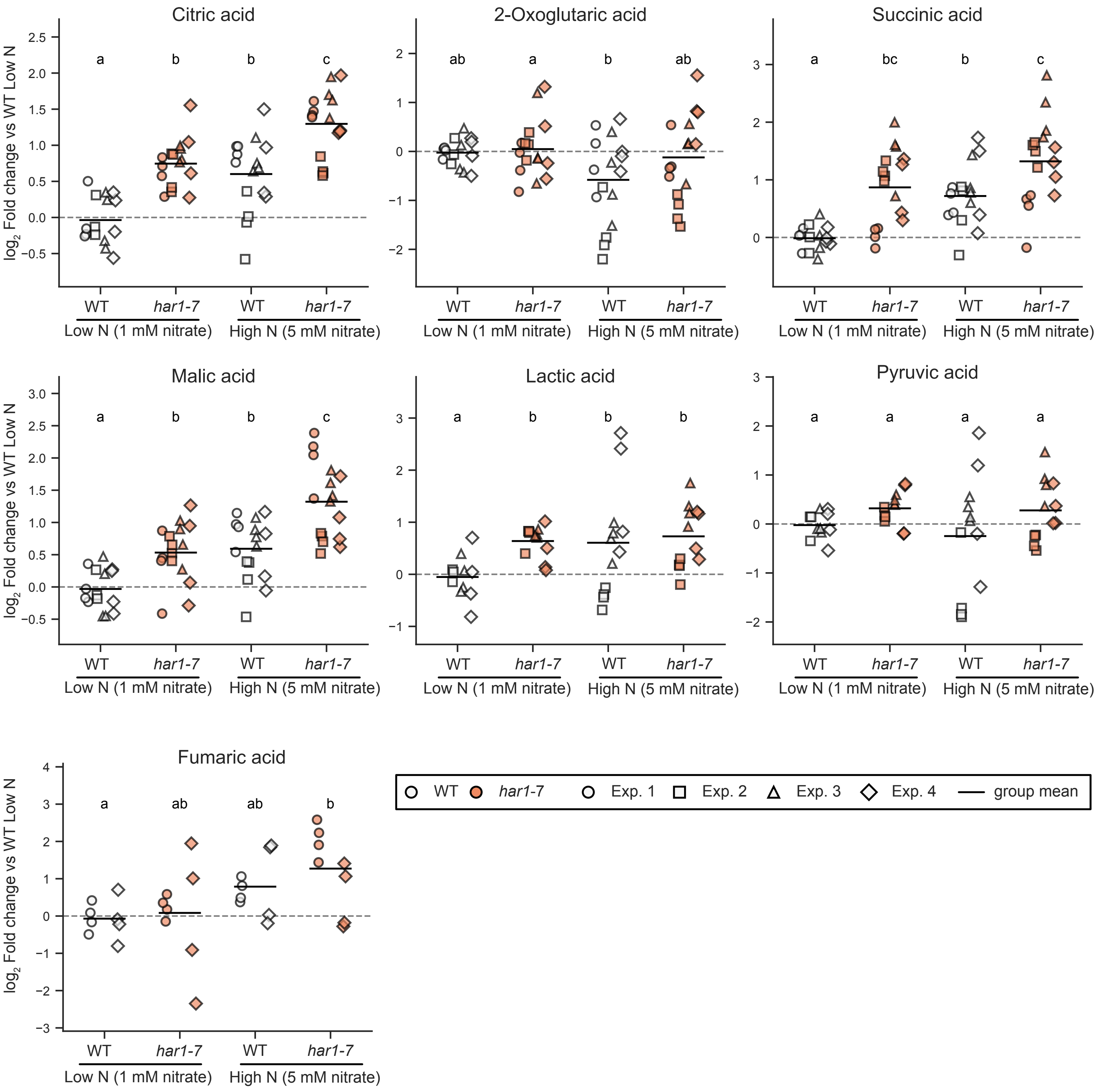


**Supplementary Figure 4. Accumulation of organic acids in wild-type and *har1-7* leaves.**
Relative levels of tricarboxylic acid (TCA) cycle-related organic acids in wild-type (WT) and *har1-7* leaves under low- and high-nitrate conditions. Values are expressed as $\log_2$ fold changes relative to the mean value of WT under low-N conditions within each experimental batch. Each dot represents an individual biological replicate; marker shapes indicate independent experimental batches. Black horizontal bars indicate group means. Different letters indicate significant differences among groups based on Tukey-adjusted pairwise comparisons of estimated marginal means from a linear model accounting for experimental batch effects ($P < 0.05$). Lactic acid and pyruvic acid were excluded from experiment 1 (Exp. 1), and fumaric acid was excluded from experiments 2 and 3 (Exp. 2 and 3), because distinct single peaks could not be reliably detected. The data for malic acid, citric acid, and succinic acid are identical to those presented in Fig. 5, but are shown here together with the other organic acids.
